# An intestinal cell type in zebrafish is the nexus for the SARS-CoV-2 receptor and the Renin-Angiotensin-Aldosterone System that contributes to COVID-19 comorbidities

**DOI:** 10.1101/2020.09.01.278366

**Authors:** John H. Postlethwait, Dylan R. Farnsworth, Adam C. Miller

**Affiliations:** Institute of Neuroscience, University of Oregon, Eugene OR 97403

**Keywords:** scRNA-seq, Apelin, COVID-19, *Danio rerio*, Angiotensinogen, conserved synteny, genome duplication

## Abstract

People with underlying conditions, including hypertension, obesity, and diabetes, are especially susceptible to negative outcomes after infection with the coronavirus SARS-CoV-2. These COVID-19 comorbidities are exacerbated by the Renin-Angiotensin-Aldosterone System (RAAS), which normally protects from rapidly dropping blood pressure or dehydration via the peptide Angiotensin II (Ang II) produced by the enzyme Ace. The Ace paralog Ace2 degrades Ang II, thus counteracting its chronic effects. Ace2 is also the SARS-CoV-2 receptor. *Ace*, the coronavirus, and COVID-19 comorbidities all regulate *Ace2*, but we don’t yet understand how. To exploit zebrafish (*Danio rerio*) as a disease model to understand mechanisms regulating the RAAS and its relationship to COVID-19 comorbidities, we must first identify zebrafish orthologs and co-orthologs of human RAAS genes, and second, understand where and when these genes are expressed in specific cells in zebrafish development. To achieve these goals, we conducted genomic analyses and investigated single cell transcriptomes. Results showed that most human RAAS genes have an ortholog in zebrafish and some have two or more co-orthologs. Results further identified a specific intestinal cell type in zebrafish larvae as the site of expression for key RAAS components, including Ace, Ace2, the coronavirus co-receptor Slc6a19, and the Angiotensin-related peptide cleaving enzymes Anpep and Enpep. Results also identified specific vascular cell subtypes as expressing Ang II receptors, *apelin*, and *apelin receptor* genes. These results identify specific genes and cell types to exploit zebrafish as a disease model for understanding the mechanisms leading to COVID-19 comorbidities.

**SUMMARY STATEMENT:** Genomic analyses identify zebrafish orthologs of the Renin-Angiotensin-Aldosterone System that contribute to COVID-19 comorbidities and single-cell transcriptomics show that they act in a specialized intestinal cell type.

## INTRODUCTION

Coronaviruses have already led to three major epidemics in the 21^st^ century: Severe Acute Respiratory syndrome caused by SARS-CoV (WHO, 2004), Middle East Respiratory Syndrome, caused by MERS-CoV (WHO, 2016), and in the worst pandemic in over a century, in December 2019, Corona Virus Disease 2019 (COVID-19) (Lu et al., 2020) caused by SARS-CoV-2, with symptoms including pneumonia, fever, persistent cough, lung inflammation, and often diarrhea, acute kidney injury, and death (Cheng Y et al., 2020; Meo et al., 2020).

Remarkably, about 20% of SARS-CoV-19 infections are asymptomatic (Bai et al., 2020; Streeck et al., 2020), which promotes surreptitious dissemination. A question of great interest is: what makes some people more susceptible to severe COVID-19 outcomes than others? Risk factors include sex, race, and age (Du et al., 2020; Jin et al., 2020; Price-Haywood et al., 2020; Richardson et al., 2020). These factors correlate with certain chronic health problems: among nearly 6000 patients hospitalized with COVID-19 in New York, 94% had a pre-existing health issue, frequently hypertension (57%), obesity (42%), and diabetes (34%) (Richardson et al., 2020). We do not yet fully understand how these underlying conditions exacerbate COVID-19, but they are related to the Renin-Angiotensin-Aldosterone System (RAAS), which modulates blood volume and blood pressure (Cabandugama et al., 2017; Fyhrquist and Saijonmaa, 2008). The RAAS is linked to COVID-19 in two important ways. First, COVID-19 comorbidities mimic a chronically over-active RAAS and second, the receptor for SARS-CoV-2 is a key RAAS component.

COVID-19 comorbidities mimic RAAS action through Angiotensin II (Ang II). This 8-amino acid peptide promotes vasoconstriction, salt and water retention, inflammation, and production of reactive oxygen species (ROS) (Supplementary Fig. S1). These features help counteract a sudden drop in blood pressure but are harmful when chronic (Fyhrquist and Saijonmaa, 2008). Some of these features contribute to COVID-19 comorbidities, including hypertension, diabetes, cardiovascular disease, and cerebrovascular disease (Wang et al., 2020). Ang II forms when Ace cleaves two amino acids from Ang I, a peptide produced by Renin digesting Angiotensinogen (Fyhrquist and Saijonmaa, 2008). Ace inhibitor drugs decrease Ang II production and thus dampen hypertension and other COVID-19 comorbidities (Rice et al., 2004) (Messerli et al., 2018; Natesh et al., 2004; Sommerstein et al., 2020). Ang II receptor blockers also improve cardiovascular health. About 19% of people hospitalized for COVID-19 had been taking one of these drugs for hypertension (Richardson et al., 2020). It is not fully known whether Ace inhibitors or Ang receptor blockers benefit or harm COVID-19 patients (Sommerstein et al., 2020; Vaduganathan et al., 2020).

COVID-19 is further linked to the RAAS because the spike protein on the surface of SARS-CoV-2 binds near the active site of Ace2, a cell-surface monocarboxy peptidase, allowing infection (Gallagher and Buchmeier, 2001; Hoffmann et al., 2020; Millet and Whittaker, 2015; Simmons et al., 2013; Walls et al., 2020; Yan et al., 2020). The normal role of Ace2 is to metabolize Ang II to Ang1-7, thus decreasing hypertension and retention of water and salt. The binding of SARS-CoV-2 to Ace2, however, inhibits Ace2 activity (Kuba et al., 2005), which would decrease the destruction of Ang II and contribute to more intense COVID-19 morbidities. Knockout mice lacking Angiotensinogen or Ang II receptors support the notion that RAAS activation leads to COVID-19 comorbidities because they show less obesity, less insulin resistance, and less hypertension (Massiera et al., 2001; Yvan-Charvet et al., 2005). Ace2 can also cleave Apelin (Apln) peptides, which cause vasodilation, increased heart muscle contractility, angiogenesis, fluid homeostasis, and regulation of energy metabolism; thus, countering effects of Ang II (De Mota et al., 2004; Dray et al., 2008; Kasai et al., 2004; Szokodi et al., 2002; Yang et al., 2017). Apln is a positive regulator of Ace2, so decreasing Apln downregulates Ace2 expression, thereby decreasing the number of virus receptors, but increasing Ang II levels, prolonging harmful effects on COVID-19 comorbidities (Sato et al., 2013).

The molecular genetic bases of complex diseases like COVID-19 are often illuminated by investigations in model organisms (Wangler et al., 2017). Among vertebrate models for human disease, zebrafish (*Danio rerio*) is a workhorse for understanding COVID-19-related organs, including heart and vasculature, kidney, liver, and many others (e.g., (Bournele and Beis, 2016; Goessling and Sadler, 2015; Morales and Wingert, 2017)). Zebrafish can even contribute to understanding communicable human diseases (Bouz and Al Hasawi, 2018; Sullivan et al., 2017); for example, transgenic cell markers and body transparency helped identify a specific macrophage type as the long-sought reservoir for asymptomatic tuberculosis (Clay et al., 2007).

Although humans are more closely related to other mammals than to fish, zebrafish can help accelerate research into the mechanisms of vertebrate health and disease. Zebrafish embryos and larvae are accessible, developing in a dish and are optically favorable, so disease mechanisms can be visualized through transgenic fluorescent imaging of functioning organs in living animals (e.g., (Wiles et al., 2016)). For example, transgenic zebrafish expressing GFP in TNFA-positive cells and mCherry in neutrophils (Lam et al., 2012; Marjoram et al., 2015) allow visualization of components of the ‘cytokine storm’ that can lead to death of adult COVID-19 patients and to the pediatric multi-system inflammatory syndrome that affects some SARS-CoV-2-infected children (Health, 2020; Ye et al., 2020; Zhang et al., 2020). Zebrafish have vertebrate-specific organs and organ systems that mimic our own and they develop and function using similar regulatory mechanisms. In addition, small size makes zebrafish suitable for screens of therapeutic molecules at a scale not possible in mammals (Lam and Peterson, 2019).

To evaluate zebrafish as a model for mechanistic insights into the links between COVID-19 comorbidities and the RAAS, we first identified the orthologs and co-orthologs of human RAAS-related genes in zebrafish. Zebrafish orthologs of human genes are sometimes obscured by genome duplication events (two rounds in stem vertebrates (Dehal and Boore, 2005; Simakov et al., 2020) and a third round in stem teleosts (Amores et al., 1998; Jaillon et al., 2004; Postlethwait et al., 1999; Taylor et al., 2001)). Second, we explored the expression of RAAS-related genes in single-cell transcriptomic (scRNA-seq) experiments from zebrafish embryos and larvae, providing the first organism-wide view of RAAS gene expression at the level of individual cell types. Analyses showed first, that the RAAS is nearly identical in zebrafish and human while identifying orthologs and previously unrecognized co-orthologs of important components; and second, that a specific under-characterized cell type expresses many RAAS components and is hence a focal cell type for the RAAS that merits further exploration. An apparently similar cell type in humans allows SARS-CoV-2 infection, the production of infectious virus, and likely some COVID-19 pathologies (Stanifer et al., 2020). These studies support zebrafish as a model for investigating the relationship of the RAAS to COVID-19 pathologies.

## RESULTS

To identify RAAS components in the zebrafish genome and to detect potential COVID-19-related cellular systems, we combined our ongoing efforts in defining the relationships of zebrafish and human genomes (Braasch et al., 2016; Braasch et al., 2015; Postlethwait et al., 1999; Postlethwait and Braasch, 2020) and our efforts in developing a scRNA-seq Atlas for zebrafish development (Farnsworth et al., 2020). Analyses described below follow an outline of RAAS function portrayed in Supplementary Figure S1 (see (Fyhrquist and Saijonmaa, 2008; Zhang et al., 2018)).

### Angiotensinogen

(Agt) is the protein precursor of Ang II (Fig. S1.1). *AGT* on human (*Homo sapiens*) chromosome 1 (Hsa1) and *agt* (ENSDARG00000016412) on zebrafish (*Danio rerio*) chromosome 13 (Dre13) conserve syntenies (Fig. 1A). Phylogenetic analysis confirmed that zebrafish has a single ortholog of human *AGT* (ENSGT00890000139531, Supplementary Fig. S2A). Mammalian and zebrafish Agt proteins share their overall 3-dimensional structure (Lu et al., 2016b) and are both strongly expressed in the adult liver (Cheng et al., 2006; Fagerberg et al., 2014)). Our zebrafish scRNA-seq Atlas, which combines cells from 1 and 2 day post fertilization (dpf) embryos and from 5dpf larvae (Farnsworth et al., 2020), extended these results, identifying two 5dpf hepatocyte sub-types (c121, c217) that strongly expressed *agt* and a third (c55) that weakly expressed *agt* (Fig. 1B, C).

**Figure 1.**
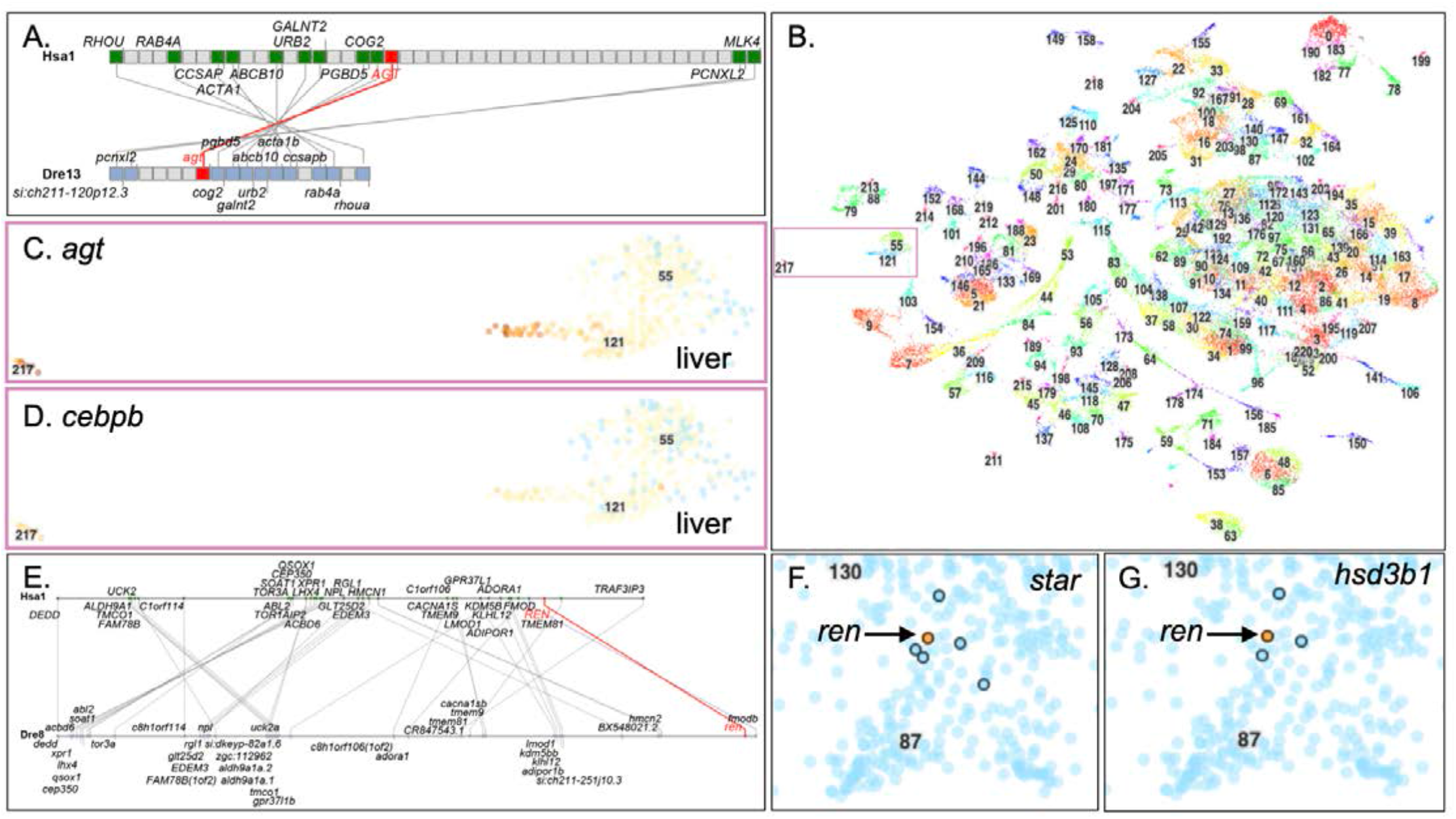
*Angiotensinogin* and *Renin*. A. The section of *Homo sapiens* chromosome 1 (Hsa1) that contains *AGT* is conserved with the segment of *Danio rerio* chromosome 13 (Dre13) that contains *agt*. B. The 220 clusters from the zebrafish scRNA-seq Atlas with liver clusters boxed (Farnsworth et al., 2020). C. Cells in hepatocyte cluster-217 (c217, left) and in a sub-set of hepatocyte cluster c121 express *agt*. Each dot represents a cell. Red color intensity indicates expression level. Blue cells are not expressing. D. Expression of the Agt-regulator *cebpb* in liver cells of 5dpf larvae. E. Conserved syntenies confirm orthology of zebrafish *ren* to *REN*. F-G. *ren* (orange cell) was expressed in c87, part of which contains precursors to the interrenal, the fish equivalent of the human adrenal cortex, as marked by the expression of the cortisol-synthesis genes *star* (F) and *hsd3b1* (G) (expressing cells circled).

Cortisol and inflammation both control *agt* expression via the glucocorticoid receptor Nr3c1 and by Cebpb, Cebpd, and other factors that bind enhancers (Brasier and Li, 1996; Demura et al., 2015). Many liver cells in the Atlas expressed *nr3c1* at low levels (Supplementary Fig. S2B,C), but other cell types, including periderm, basal skin cells, ionocytes, fast skeletal muscle, photoreceptors, and vascular endothelium, expressed *nr3c1* at higher levels. Nearly all *agt*-expressing cells in 5dpf zebrafish livers expressed the *agt*-regulator *cebpb* in the Atlas, while liver cells not expressing *agt* generally didn’t express *cebpb* (Fig. 1D) or *cebpd* (Supplementary Fig. S2D). This result shows that genes expected to initiate the RAAS are already in place in hepatocyte cell types in 5dpf zebrafish.

### Renin

encodes the enzyme that cleaves Ang I from Angiotensinogen (Fig. S1.1). Conserved syntenies (Fig. 1E) support the orthology of human *REN* to zebrafish *ren* (ENSDARG00000041858) (Liang et al., 2004). Phylogenetics (ENSGT00940000157898, Fig. S3A) confirmed a single copy of *ren* in zebrafish. *Renin* is expressed by kidney juxtaglomerular cells in adult mammals and adult zebrafish (Gomez et al., 1988; Hoshijima and Hirose, 2007; Jones et al., 1990; Liang et al., 2004). In mammalian embryos and fetuses, however, *Ren*-expressing cells arise predominantly from the adrenal but also other primordia, including skin, nervous system, spleen, testis, eyes, and others (Gomez et al., 1988; Jones et al., 2000; Sequeira Lopez et al., 2004). Only three cells in the Atlas expressed detectable levels of *renin*, and, as in mammalian fetuses, each of the zebrafish *ren* cells was in a different cluster. As predicted from mammals and zebrafish (Gomez et al., 1988; Liang et al., 2004; Sequeira Lopez et al., 2004), one of these cells was an interrenal (adrenal homolog) precursor (c89) verified by expression of steroidogenic genes *star* and *hsd3b1* (Fig. 1F,G). The hypothesis that *Renin, Ctsd*, and *Napsa* derived by duplication events in stem bony vertebrates (Liang et al., 2004) derived support from our genomic analyses showing that the portions of Hsa1, Hsa11, and Hsa19 that contain these three genes, along with a part of Hsa12, make up four ohnologous regions (Supplementary Figure S3B) predicted by two rounds of Vertebrate Genome Duplication (VGD (Dehal and Boore, 2005; Simakov et al., 2020)).

### Angiotensin I

(Ang I, Fig. S1.1) varies in sequence among vertebrates (Takei et al., 1993). To follow the evolution of Ang I sequences, we downloaded Agt sequences across vertebrate phylogeny. Most placental mammals have the same Ang I sequence as humans, likely the ancestral state for eutherian mammals (DRVYIHPFHL, Fig. 2). Some Artiodactyls, like cows and whales, however, have a valine (V) at position 5, as do squirrel-related mammals and some bats. These replacements are due to lineage-specific mutations because each is embedded in a clade with isoleucine at position 5. The little brown bat *Myotis lucifugus* has the most variant Ang I among sampled therian mammals, with leucine rather than the otherwise invariant valine at position 3 and methionine rather than the otherwise invariant leucine at position 10. This Ang 1-divergent bat belongs to the group (Vespertilioniodea) that harbors coronaviruses most closely related to SARS-CoV-2 and are the presumed wild source of this zoonotic virus (Anthony et al., 2017; Cotten et al., 2013; Lau et al., 2020; Toosy AH and S., 2019). Monotremes and marsupials have valine rather than isoleucine at position 5, which appears to be the ancestral mammalian state (DRVYVHPFHL) because most non-mammalian tetrapods also have valine at this position. Most non-mammalian sarcopterygians (lobe-finned vertebrates) have, like mammals, aspartic acid at position 1, although basally diverging lobe-finned vertebrates, including amphibia and coelacanth, have asparagine at this position. Because actinopterygians (ray-finned vertebrates) also have asparagine at position 1, this is likely the ancestral state for all osteichthys (bony vertebrates). Although mammals have histidine at position 9, non-mammalian lobe-finned vertebrates have a variety of amino acid residues at this position. Because the most basally diverging lobe-finned vertebrates (clawed frog *Xenopus tropicalis* and coelacanth) both have asparagine at position 9, and because basally diverging ray-finned vertebrates also have asparagine at position 9, the sequence NRVYVHPFNL is likely the ancestral state for lobe-finned vertebrates.

**Figure 2.**
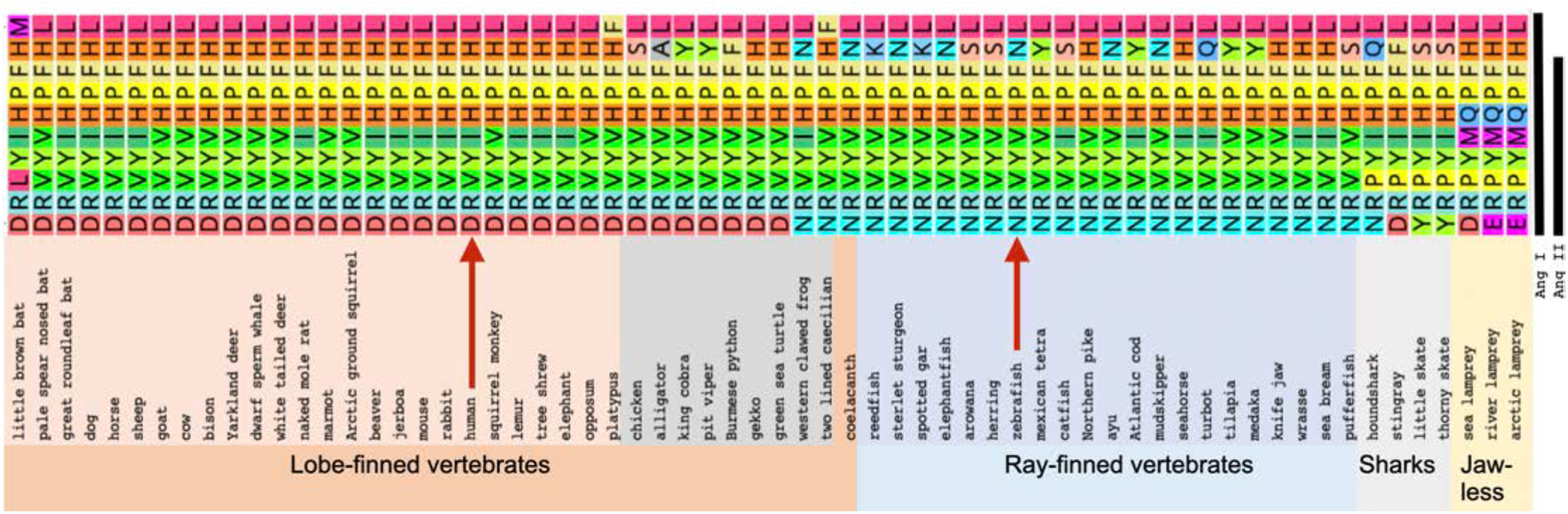
Evolution of Angiotensin sequences. Arrows indicate human and zebrafish sequences. Species organized according to published phylogenies (Hughes et al., 2018; Upham et al., 2019). The extent of Ang I and Ang II indicated at far right. For Latin names and accession numbers, see Supplementary Table S1.

Ang I sequences vary more among ray-finned than lobe-finned vertebrates (Fig. 2). Because coelacanth and several basally diverging ray-finned vertebrates (sturgeon, elephant fish) have the sequence NRVYVHPFNL, this is likely the ancestral Ang I sequence for all bony vertebrates. Several ray-finned vertebrates have isoleucine at position 5, representing independent mutations that happen to match placental mammals. Position 9 is highly variable in ray fins as in lobe fins. Zebrafish Ang I is NRVYVHPFNL, differing from the human form at positions 1, 5, and 9.

The Angiotensin system was likely already active in stem vertebrates because chondrichthys (cartilaginous fish) and even agnathans (jawless vertebrates) possess Angiotensinogen genes (Takei et al., 1993). At position 1, some cartilaginous fish have arginine like most ray-finned vertebrates, but others have asparagine or tryptophan, and position 9 is variable. The ancestral Ang I sequence in jawed vertebrates was likely the same as in ancestral bony fishes (NRVYVHPFNL).

Angiotensinogen genes in jawless vertebrates appear to encode an Angiotensin that shares the amino-terminal four residues with mammals but varies in the carboxy-terminal six residues (Wong and Takei, 2011). At position 1, lampreys have either aspartic acid or glutamic acid, but the conserved isoleucine or leucine at position 5 is replaced with methionine followed by glutamine replacing the otherwise invariant histidine.

Lamprey Ang II alters cardiovascular dynamics in live lampreys but teleost Ang II (NRVYVHPF) did not (Wong and Takei, 2011), showing that stem jawless vertebrates already had key components of the RAAS but that ligands and receptors evolved differences. Atg searches against the non-vertebrate chordate amphioxus returned serine protease inhibitors (SERPINs), but not Angiotensinogin, suggesting that Ang I signaling originated in vertebrates after divergence from non-vertebrate chordate ancestors.

### Angiotensin converting enzyme

(Ace) changes Ang I to Ang II (Fig. S1.2). Syntenies are conserved between *ace* (ENSDARG00000079166) in zebrafish and *ACE* (ENSG00000159640) in human supporting orthology (Fig. 3A). In humans, *ACE* is expressed four times stronger in the small intestine than in lungs (Fagerberg et al., 2014) and in 5dpf zebrafish, *ace* is also expressed in the gut (Rauch et al., 2003). In the Atlas *ace* was expressed almost exclusively in the 5dpf intestinal epithelium cluster c152 (Fig. 3B-D). The most statistically differentially expressed genes in the *ace-*expressing cluster c152 encode fatty acid binding proteins and apolipoproteins, suggesting a function in lipid biology.

**Figure 3.**
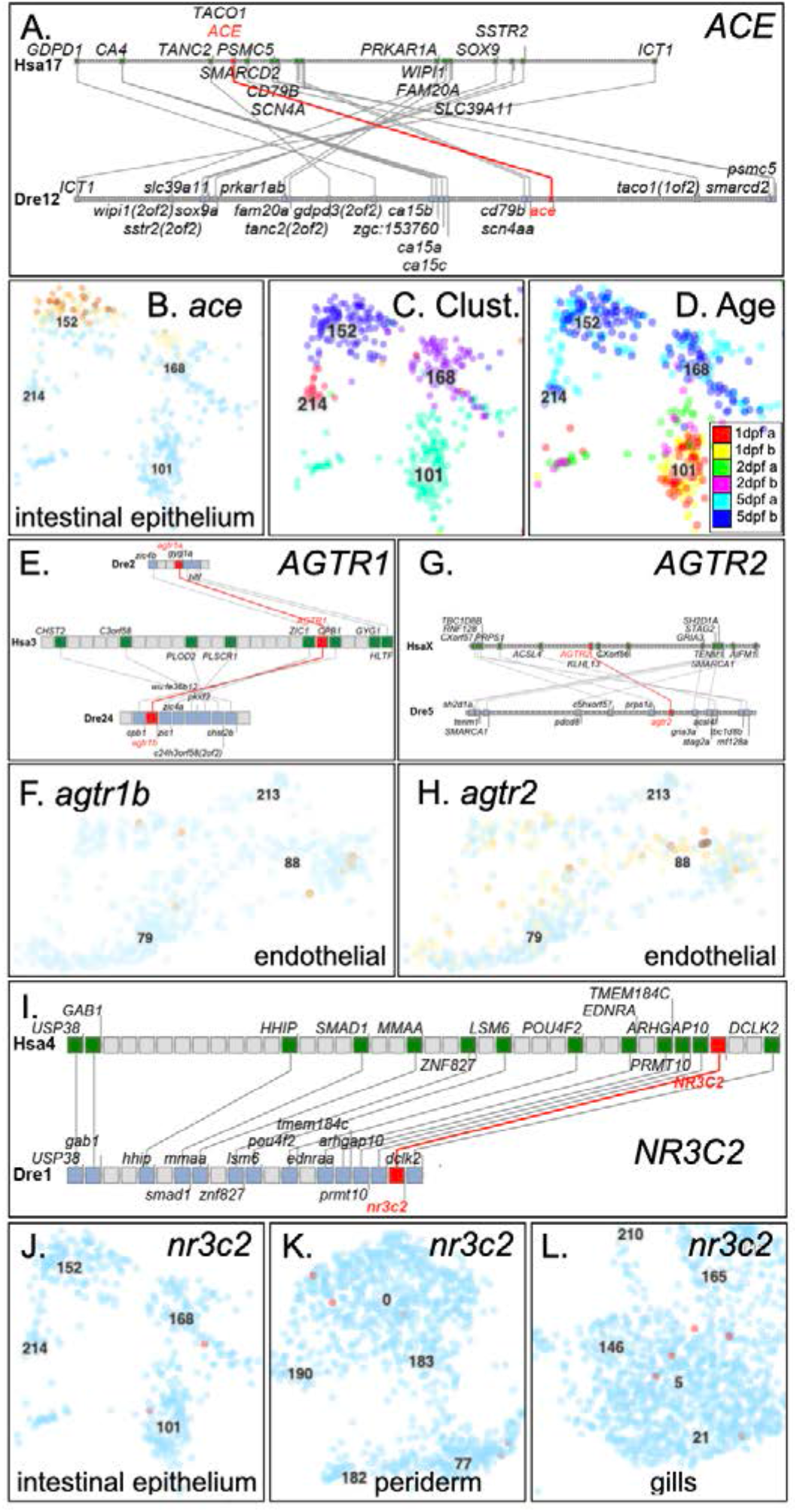
Genomics and expression of *ace, agtr*, and *nr3c2*. A. Conserved syntenies verify orthology of zebrafish *ace* to human *ACE*. B. Expression of *ace* in a specific intestinal epithelial cell type in c152. C. Clusters c101, c152, c168, and c214 are intestinal epithelial cell types (Farnsworth et al., 2020). D. The age of each cell, as indicated by colors displayed in the insert, showed that *ace* is expressed in a 5dpf intestinal cell type that does not exist at 1dpf or 2dpf (c101). E. Double conserved synteny of zebrafish *agtr1a* and *agtr1b* to human *AGTR1*. F. *agtr1b* was expressed only in endothelial cells (c88). Expression of *agtr1a* was not detected. G. Conserved synteny of *agtr2* with *AGTR2*. H. Expression of *agtr2* was detected in c88 endothelial cells, including some in which *agtr1b* was detected. I. Conserved syntenies for the aldosterone receptor gene *nr3c2*. J. Expression detected for *nr3c2* in intestinal epithelial clusters c168 and c101. K.-L. Expression of *nr3c2* in the periderm at 1 and 2dpf and in the gills at 5dpf.

### Angiotensin II

(Ang II) forms when Ace cleaves two C-terminal amino acid residues from Ang I (Fig. S1.3). Ang II contributes to hypertension, an important COVID-19 comorbidity, and likely promotes the inflammation that leads to poor disease outcomes. Human and zebrafish Ang II differ only at the first and fifth residues (DRVTIHPF vs. NRVTVHPF, Fig. 2). Importantly, the fish and human peptides act about equally in stimulating heterologous systems (Russell et al., 2001; Vilella et al., 1996). Treating zebrafish with Ang II causes increased sodium uptake as expected if Ang II function is conserved between human and zebrafish (Kumai et al., 2014).

### Angiotensin II receptor type 1 and type 2

(Agtr1, Agtr2, Fig. S1.4) transduce Ang II signals on vascular and other cells. In mammals, Agtr1 mediates inflammation and the production of aldosterone in the adrenal cortex, thereby increasing the kidney’s reabsorption of sodium (Higuchi et al., 2007). Agtr2 broadly opposes Agtr1 by decreasing cell proliferation and vasoconstriction (Wang et al., 1998b). Zebrafish has two co-orthologs of *AGTR1* (*agtr1a* (ENSDARG00000018616) and *agtr1b* (ENSDARG00000045443)) that reside in double conserved syntenies with human *AGTR1*, verifying orthology (Fig. 3E). (The gene formerly called *agtrl1a* (Tucker et al., 2007) is now recognized as an Apelin receptor gene *aplnra* (ENSDARG00000002172)). Human *AGTR1* and *AGTR2* are expressed in vascular endothelial cells; likewise, the Atlas showed *agtr1b* expression strongest in endothelial cells (c88, Fig. 3F) and in an unidentified type of mesenchyme (c135), likely vascular precursors. In mammals, the kidney controls ion and water balance and expresses *Agtr1* (Miyata et al., 1999), but in fish, ionocytes in the skin control ion and water balance (Dymowska et al., 2012; Guh et al., 2015; Inokuchi et al., 2017; Kwong et al., 2016) although they did not have detectable *agtr1a* or *agtr1b* expression in the Atlas despite the fact that eel ionocytes express a protein that binds a mouse antibody to Agtr1 (Marsigliante et al., 1997). Expression of *agtr1a* was not detected in the Atlas.

Zebrafish has a single ortholog of human *AGTR2* with conserved syntenies (Fig. 3G) (Wong and Takei, 2013). *In situ* hybridization identified *agtr2* (ENSDARG00000035552) expression in zebrafish endothelia (Wong et al., 2009), which the Atlas confirmed (Fig. 3H). Expression of *agtr2* was also detected in 5dpf mesenchyme (c135) and in 5dpf enteric smooth muscle cells (c197). Agtr antagonists taken to control hypertension also ameliorate zebrafish models of heart failure (Quan et al., 2020), showing conserved structure and function of angiotensin receptors.

In mammals, Agtr1 mediates adrenal secretion of aldosterone (Rainey et al., 2004). Teleosts have no aldosterone, lacking an ortholog of the aldosterone-synthesizing enzyme *Cyp11b2*, but nevertheless have an ortholog of *Nr3c2*, which encodes the aldosterone (mineralcorticoid) receptor (Bridgham et al., 2006). In fish, Nr3c2 is stimulated either by the aldosterone precursor 11-deoxycorticosterone or by cortisol (Kumai et al., 2014; Sturm et al., 2005). Human and zebrafish share strong conserved syntenies around *NR3C2* and *nr3c2* (ENSDARG00000102082, Fig. 3I). In the Atlas, *nr3c2* was expressed in a few cells of the 5dpf intestinal epithelium (c101, c168), in the periderm in 1dpf and 2dpf embryos (c0), in the gills in 5dpf larvae (c5) (Fig. 3J-L), and in one NaK ionocyte cell (c128), cell types that contribute to water and salt balance at these stages (Dymowska et al., 2012; Fu et al., 2010; Hoffmann et al., 2018; Rombough, 2007). Zebrafish embryos also expressed *nr3c2* in some retinal progenitors and a few other scattered cells in the Atlas.

### Angiotensin I converting enzyme-2

(Ace2, Fig. s1.7) is the SARS-CoV-19 receptor (Hoffmann et al., 2020) but its normal function is to remove an amino acid from Ang II to form Ang 1-7 (the seven amino terminal residues in Fig. 2). Phylogenetics (ENSGT00940000158077, Fig. S4A) and conserved syntenies (Fig. 4A) support orthology of human *ACE2* (ENSG00000130234) to zebrafish *ace2* (ENSDARG00000016918) (Chou et al., 2006). Human *ACE2* is expressed 3-to 4-times stronger in the intestine than in gall bladder, kidney, and testis, 6-fold stronger than in heart, and 190-fold stronger than in lungs (Fagerberg et al., 2014). *ACE2* is also strongly expressed by epithelial cells of the human oral mucosa (Xu et al., 2020), perhaps providing a route of infection. In the Atlas, *ace2* was expressed exclusively in c152, the same intestinal epithelial cell cluster that expressed *ace* (Fig. 4B vs. 3B). The co-expression of *ace* and *ace2* identifies this incompletely characterized cell type of the digestive tract as a key mediator of RAAS activity and may help account for the gastrointestinal pathologies of COVID-19 that often occur in children and adults (Guan et al., 2020; Health, 2020).

**Figure 4.**
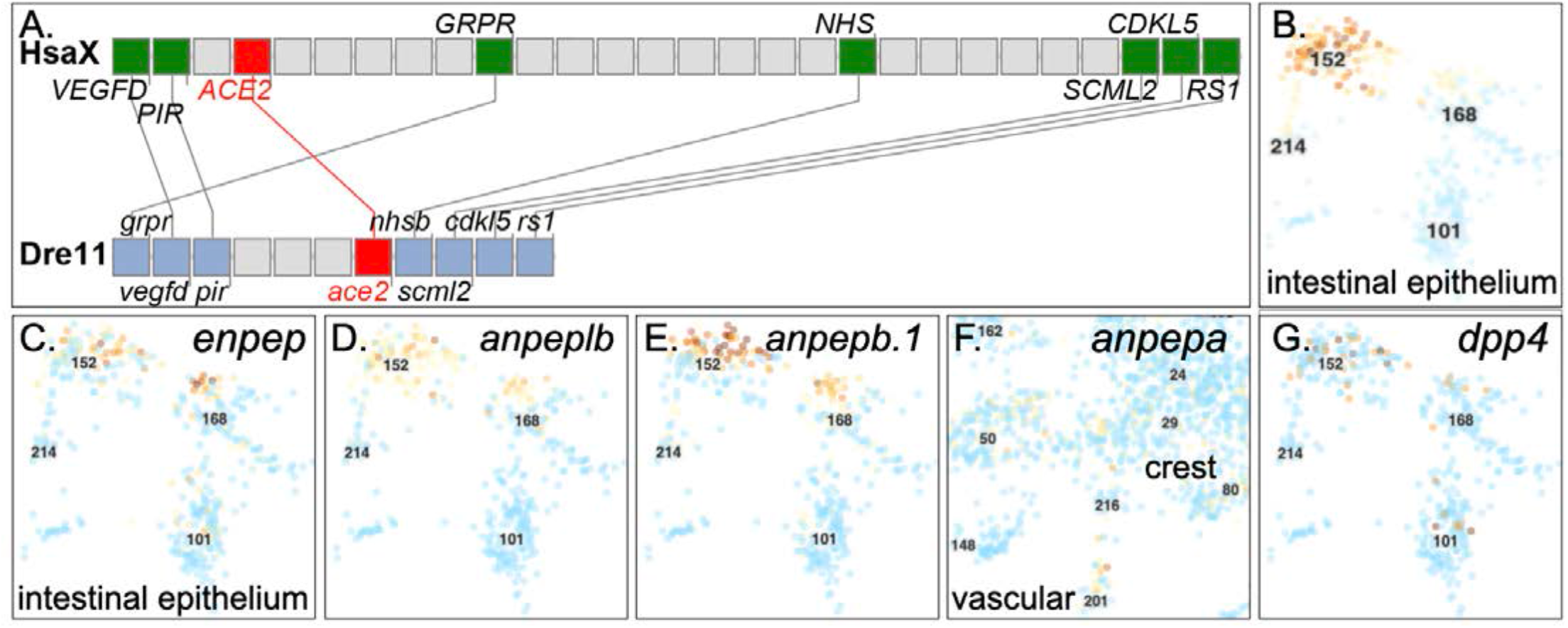
Genomics and expression of *ace2, enpep, anpep* and *dpp4*. A. Conserved synteny supports the orthology of human *ACE2* to zebrafish *ace2*. Expression of *ace2* (B), *enpep* (C), *anpeplb* (D), *anpepb*.*1*, (E), *anpepa* (F), and *dpp4* (G). All were expressed in c152, the same intestinal epithelium cell type that expressed *ace*, except *anpeplb*, which was expressed in blood vessels at 5dpf and a few neural crest cells, but not in intestinal epithelium.

### Mas1

is a receptor for Ang 1-7 (Fig. S1.8,9) (Santos et al., 2003). *MAS1* is present in mammals, sauropsids, and amphibians, and a related gene (ENSLACG00000015240) is found in coelacanth, a basally diverging lobe-finned vertebrate, but appears to be missing from ray-finned vertebrates (Fournier et al., 2012). Conserved syntenies suggest a mechanism for this situation. In human, *MAS1* lies in the gene sequence: *TCP-MRPL18-MAS1-IGF2R-SLC22A1-SLC22A2*. Zebrafish orthologs of genes to the right of *MAS1* are adjacent as expected, but zebrafish orthologs of genes to the left of *MAS1* are on different chromosomes in both zebrafish and spotted gar, a basally diverging ray-finned vertebrate (Braasch et al., 2016). Thus, the genomic neighborhood expected to contain the ray-fin ortholog of *MAS1* is rearranged with respect to the human genome, consistent with either gene loss due to a chromosome rearrangement breakpoint in stem ray-fins, or to the origin of *MAS1* in stem lobe-finned vertebrates. A different G-protein coupled receptor (GPCR) in this large protein family (Rinne et al., 2019) may act as a zebrafish Ang 1-7 receptor.

### Enpep

(glutamyl aminopeptidase) converts Ang II to Ang III (Fig. S1.10), thereby stimulating systemic blood pressure (Mizutani et al., 2008). Zebrafish has a single *enpep* (ENSDARG00000057064) gene on Dre13; phylogenetic analysis (ENSGT00940000156946) and conserved syntenies with human *ENPEP* on Hsa4 (Fig. S4B,C) verify orthology. Cells in the Atlas that expressed *enpep* also expressed *ace* and *ace2* in c152 and c168 (Fig. 4C vs. Figs. 3B, 4B), increasing the relevance of this cell type to COVID-19.

### Anpep

(alanyl aminopeptidase) metabolizes Ang III to Ang IV (Fig. S1.11) and is a receptor for the human common cold coronavirus HCoV-229E (Fehr and Perlman, 2015). Humans have a single *ANPEP* gene, but zebrafish has five *ANPEP*-related genes distributed on three chromosomes. To connect the zebrafish genome to human biology, we analyzed gene trees and conserved syntenies. Results showed that before the divergence of lobe-finned and ray-finned vertebrates, a tandem duplication produced *anpep* and *anpepl* genes (Fig. S5,S6). The *anpepl* gene was lost in stem lobe-finned vertebrates. In ray-finned vertebrates, the TGD produced ‘*a*’ copies of both *anpep* and *anpepl* on one chromosome and ‘*b*’ copies of both genes on the duplicate chromosome. One chromosome became Dre7 containing *anpepa* (ENSDARG00000036809, ZFIN *si:ch211-106j24*.*1*) and *anpepla* (ENSDARG00000089706, ZFIN *si:ch211-276a23*.*5*). The ‘b’ copy fates were more complex: first, a previously identified chromosome fission event (Nakatani and McLysaght, 2017) separated *anpeplb* (ENSDARG00000041083, ZFIN: *anpepa*) on Dre18 from the temporary *anpepla* gene on Dre25, which then duplicated to form the tandemly duplicated *anpeplb*.*1* (ENSDARG00000103878, ZFIN: *anpepb*) and *anpeplb*.*2* (ENSDARG00000097285, ZFIN: *si:ch211-147g22*.*5*). This genomic analysis clarifies relationships of zebrafish *ANPEP* paralogs necessary to connect human and zebrafish biology.

In human, ANPEP is expressed about six times stronger in the duodenum and small intestine than in the third highest organ, the kidney (Fagerberg et al., 2014). In zebrafish, *anpepb* and *anpeplb*.*1* were expressed in the same intestinal cell type as *ace, ace2*, and *enpep* (c152) (Fig. 4D, E). In contrast, *anpepa* was not expressed in the intestine, but was expressed in a 5dpf blood vessel cell type (c201, Fig. 4F) different from the cell types expressing Angiotensin receptors *agtr1b* and *agtr2*, as well as in a few cranial neural crest cells that had developed by 5dpf. The other two *ANPEP*-related zebrafish genes (*anpepla, anpeplb*.*2*) were not expressed in the Atlas. These five zebrafish *ANPEP-*related genes may share among them the original functions of the ancestral *anpep* gene, some or most of which might have been retained by the human *ANPEP* gene.

### DPP4

(dipeptidyl peptidase-4) is the receptor for MERS-CoV (Raj et al., 2013). DPP4 inhibitors also inhibit Ace and are anti-diabetic (Abouelkheir and El-Metwally, 2019). Ang II stimulates DPP4 activity in the mammalian kidney (Aroor et al., 2016). Zebrafish *dpp4* (ENSDARG00000079420) shares syntenies with human *DPP4*, verifying orthology (Fig. S7). In the zebrafish Atlas, *dpp4* was expressed exclusively in the same cells that express *ace, ace2, enpep, anpepa*, and *anpepb* (Fig. 4G), improving the resolution of *in situ* hybridization (Thisse and Thisse, 2004). These results show that zebrafish orthologs of receptors for at least three coronaviruses (SARS-CoV-2 (Ace2), HCoV-229E (Anpep), MERS-CoV (Dpp4)) are expressed almost exclusively in the same specialized cell type of the larval digestive tract.

### Slc6a19

(Fig. S1.12) is a neutral amino acid transporter and forms a heteromeric dimer with ACE2, the human SARS-CoV-19 receptor (Yan et al., 2020). Zebrafish has three *SLC6A19* co-orthologs: two (*slc6a19a*.*1* (ENSDARG00000018621) and *slc6a19*.*2* (ENSDARG00000091560)) are tandem duplicates on chromosome Dre19 and the third (*slc6a19b*, ENSDARG00000056719) on Dre16 represents the TGD duplicate (Fig. 5A), confirmed by the phylogenetic tree (Fig. S8). Double conserved synteny of *slc6a19* paralogs on Dre19 and Dre16 with human *SLC6A19* (ENSG00000174358) on Hsa5 confirms orthology (Fig. 5A). *SLC6A19* expression is about equally in small intestine, duodenum, and kidney, about 22-fold stronger than the next organ, the colon (Fagerberg et al., 2014). In the Atlas, all three *slc6a19* paralogs were expressed exclusively in the same intestinal cell type as *ace2* (c152, Fig. 5B-D), as expected if Slc6a19 proteins interact in a similar way in zebrafish and human.

**Figure 5.**
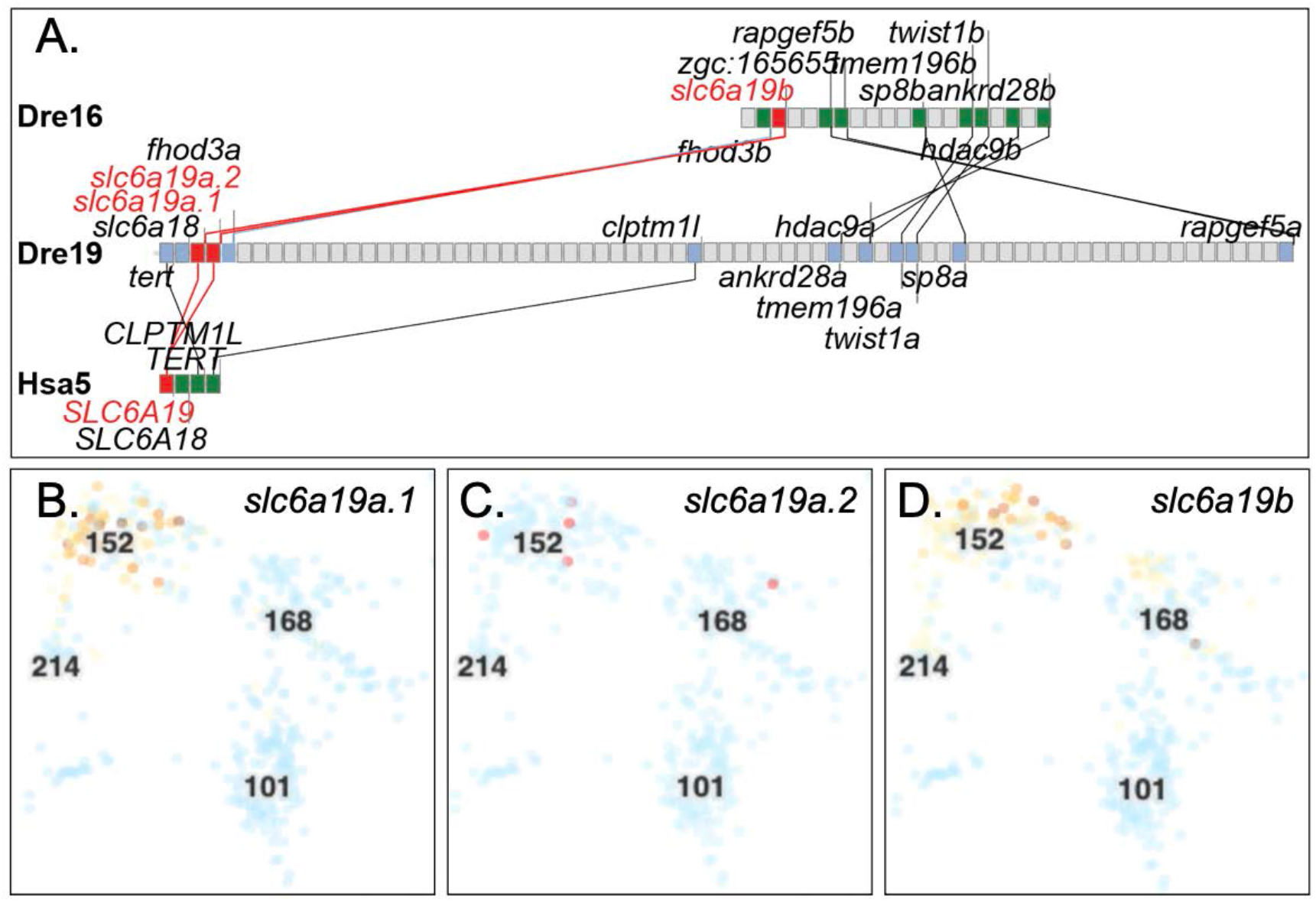
Conserved syntenies and scRNA-seq expression of zebrafish orthologs of *SLC6A19*. A. Conserved syntenies showing paralogy of a part of Dre16 containing *slc6a19b* and a portion of Dre19 containing *slc6a19a*.*1* and *slc6a19*.*2* and their relationship to human chromosome Hsa5 around *SLC6A19*. B-D. Expression of *slc6a19a*.*1, slc6a19a*.*2*, and *slc6a19b*, respectively in the zebrafish Atlas was detected in c152 and c168, which represent the same larval intestinal epithelial cells that were expressing *ace* and *ace2*.

The role of Slc6a19 in COVID-19 is as yet unknown. Hints, however, come from recent population genetic studies. *SLC6A19* and *SLC6A18* paralogs are adjacent in human and zebrafish and, along with *SLC6a20* on Hsa3, represent a closely related phylogenetic gene clade (Kristensen et al., 2011). In birds, *Slc6a19* (ENSJHYG00000013510) and *Slc6a18* (ENSJHYG00000013530) loci are adjacent and syntenic with *Slc6a20* (ENSJHYG00000002729). The phylogeny and genomics imply origin by tandem duplication followed by translocation in mammals after their divergence from the bird lineage. Like *Slc6A19* and *Ace2, Slc6A20* is expressed in intestinal enterocytes and kidney cells (Romeo et al., 2006). Furthermore, SLC6A20 interacts functionally with ACE2 (Vuille-dit-Bille et al., 2015). Importantly, recent analyses showed that *SLC6A20* is at the peak of a genome wide association study for poor outcomes from COVID-19 (Ellinghaus et al., 2020).

These findings raise the novel hypothesis that SLC6A20 variants contribute to variation in COVID-19 outcomes due to differences in expression or protein function related to interactions with ACE2, the SARS-CoV-2 receptor.

### Adam17

(Fig. S1.13) is a metalloendopeptidase that cleaves the membrane isoform of Ace2 to make the soluble protein sAce2 (Lambert et al., 2005). Zebrafish has two copies of *adam17* in double conserved synteny with human *ADAM17* (Fig. 6A), showing that they are co-orthologs of human *ADAM17* from the TGD.

**Figure 6.**
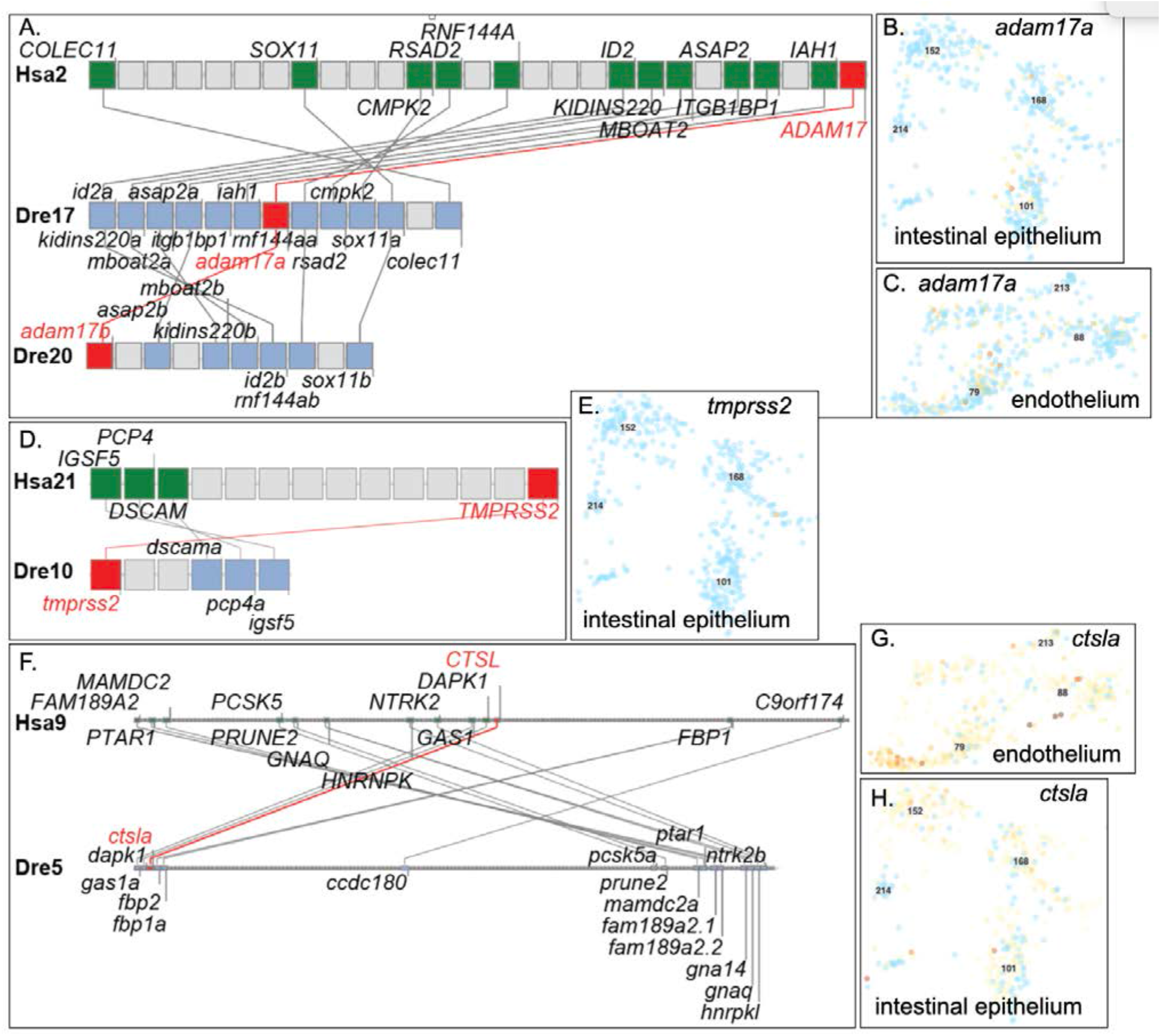
Conserved syntenies and expression of a*dam17, tmprss2*, and c*tsl*. A. Double conserved synteny between Dre17 and Dre20 confirming orthology of zebrafish *adam17a* and *adam17b* to human *ADAM17* and their origin in the teleost genome duplication. B. Expression of *adam17a* weakly in intestinal epithelium cells and (C.) stronger in the vascular endothelium. D. Conserved syntenies verify orthology of *tmprss2* to *TMPRSS2*. E. Expression of *tmprss2* was detected in one cell in intestinal epithelium c168 and in 12 other cells broadly dispersed in the atlas. F. Human *CTSL* shares conserved syntenies with zebrafish *ctsla*. G, H. Expression of *ctsla* in the endothelium and intestinal epithelium, respectively.

The zebrafish Atlas showed expression of *adam17a* (ENSDARG00000043213) stronger in embryonic intestinal epithelium than in larval *ace2*-expressing c152 cells (Fig. 6B). In addition, several cells in the vascular endothelium expressed *adam17a* (Fig. 6C). The TGD duplicate *adam17b* (ENSDARG00000093093) was expressed in the Atlas in a few widely dispersed individual cells. Because expression of *adam17* paralogs was not detected in c152, zebrafish may not have soluble Ace2; alternatively, a different enzyme in zebrafish might cleave Ace2 to make the soluble form or expression levels may be too low for detection.

### Tmprss2

(Fig. S1.16) is a transmembrane serine protease that activates the coronavirus spike protein, stimulating virus entry into cells (Hoffmann et al., 2020; Millet and Whittaker, 2015; Walls et al., 2020). In humans, *TMPRSS2* expression is highest in prostate, followed by the digestive system, and below that, the lung (Fagerberg et al., 2014). *TMPRSS2* in human and *tmprss2* (ENSDARG00000098686) in zebrafish occupy regions of conserved synteny, confirming orthology (Fig. 6D). Only one intestinal epithelial cell expressed *tmprss2* in the Atlas (c168, Fig. 6E) as well as three kidney cells (not shown).

### Ctsl

(cathepsin L) can cleave the spike protein of SARS-CoV (Simmons et al., 2005). *CTSL* on Hsa9 shows strong conserved synteny with zebrafish *ctsla* (ENSDARG00000007836) on Dre5 (Fig. 6F). Zebrafish *ctslb* (ENSDARG00000074306) on Dre12 is in a cluster of 12 *ctsl-*related tandemly duplicated paralogs; it does not show conserved synteny with human *CTSL* and thus does not support the conclusion that *ctslb* is a TGD paralog of *ctsla*. In the Atlas, *ctslb* is expressed exclusively in the hatching gland, the egg-shell digesting organ (Rauch et al., 2003 ; Thisse and Thisse, 2004). In contrast, *ctsla* is expressed in the endothelium (Fig. 6G), along with *agtr1b* and *agtr2* (Fig. 3F, H), in the intestine, like *ace* and *ace2*, and in liver, kidney, spleen, and gill. Because Ctsl cleaves the spike protein of SARS-CoV (Simmons et al., 2005), it may also activate the spike protein of SARS-CoV-2; expression of *ctsla* in zebrafish *ace2-*expressing cells makes it a likely candidate to model SARS-CoV-2 function in COVID-19.

### Apelin

(Apln, Fig. S1.17-18) peptidets oppose the effects of Ang II (De Mota et al., 2004; Dray et al., 2008; Kasai et al., 2004; Szokodi et al., 2002). Zebrafish has a single copy of *apln* (ENSDARG00000053279, see ENSGT00390000014020) (Quertermous, 2007; Scott et al., 2007; Zeng et al., 2007) that has strong conserved synteny with human (Fig. S9). Expression of *apln* in zebrafish has been reported in the axial chordamesoderm, notochord, heart primordium, and vasculature (Helker et al., 2015; Kwon et al., 2016; Scott et al., 2007; Zeng et al., 2007). The Atlas confirms these expression domains; for example, *apln* was expressed in endothelial cell type c88 (Fig. 7E) at 1,2 and 5dpf but not in the other two endothelial cell types. The co-expression of venous markers (*efnb2a, notch3, hey2*, and *tbx20*) and arterial markers (*flt4, ephb4*) (Fischer et al., 2004; Lawson et al., 2001; Saint-Geniez et al., 2003; Szeto et al., 2002; Thompson et al., 1998; Wang et al., 1998a; Zhou et al., 2015) show that *apln* was expressed in venous endothelium. In addition, *apln* was expressed in cardiac muscle (c205) at 1 and 2dpf, cardiac neural crest (c69) at 1dpf, vessel precursors (c117) at 1dpf, NCC type ionocytes at 1 and 2dpf, photoreceptors (c115) at 5dpf, and notochord (c158, c149). These results confirm and extend prior studies (Helker et al., 2015; Kwon et al., 2016; Scott et al., 2007; Zeng et al., 2007). A zebrafish knockout mutant allele had normal vasculogenesis (Helker et al., 2015), suggesting that Apln is not required for development of the vessels that secrete it, but affects regulates the migration of myocardial progenitors to the anterior lateral plate mesoderm (Helker et al., 2015; Scott et al., 2007; Zeng et al., 2007).

**Figure 7.**
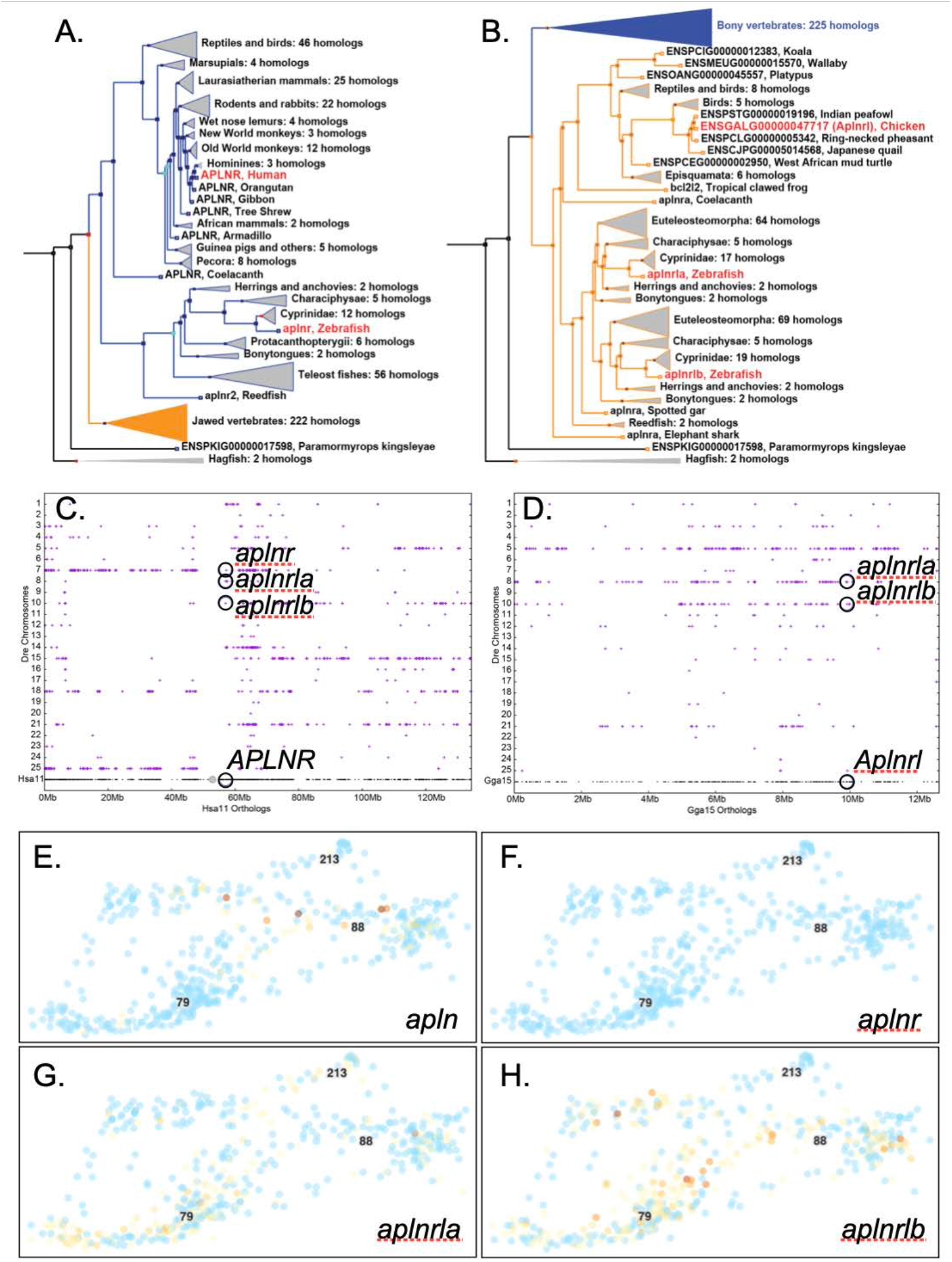
Apelin and its receptors. A. Phylogenetic tree ENSGT01000000214406 with the *APLNR/aplnr2* (or *aplnr* (ENSDARG00000004447) subtree (blue) expanded. B. The same tree with the *aplnra/aplnrb* (or *aplnrla*, ENSDARG00000002172) / *aplnrlb* (ENSDARG00000036670)) subtree (orange) expanded. C. A dot plot representing orthologs and paralogs of Hsa11 genes vs. zebrafish (Dre) chromosomes plotted directly above the location of Hsa11 genes showing extensive conservation with Dre7, the site of *aplnr2*, but not Dre8 or Dre10, the locations of *aplnra* and *aplnrb*, respectively. D. A dot plot of chicken chromosome Gga15 with vs. zebrafish chromosomes showing the sites of *aplnra* and *aplnrb* and their ortholog in chicken (ENSGALG00000047717) identified in the tree in panel B. E.-H. Expression of *apln, aplnr2, aplnra*, and *aplnrb* in endothelial cells.

### Apln receptor

(Aplnr, alias Agtr1l1 or Apj) is single copy in mammals but has three copies in zebrafish (*aplnra* (ENSDARG00000002172), *aplnrb* (ENSDARG00000036670), and *aplnr2* (ENSDARG00000004447, alias *aplnr3a* (Zhang et al., 2018))). It is claimed (Zhang et al., 2018) that these names do not reflect mammalian orthologs. To resolve this controversy, we investigated phylogenetic trees and conserved syntenies. Phylogenetic analysis for *APLNR*-related genes (ENSGT01000000214406) showed that human *ALPNR* (ENSG00000134817) branched with chicken *Alpnr* (ENSGALG00000004841) and zebrafish *aplnr2* (ENSDARG00000004447), as expected if these three genes were orthologs (Fig. 7A). Analysis of conserved syntenies showed that about half of Hsa11, including Hsa11p and the proximal part of Hsa11q that contains *ALPNR*, are orthologous to the portion of Dre7 that contains *aplnr2* (Fig. 7C). The TGD copy of this part of Hsa11 includes parts of Dre25 and Dre14 or Dre21, which lost the co-ortholog of *aplnr2* (Fig. 7C). Dre8 and Dre10, the locations of *aplnra* and *aplnrb*, respectively, had substantially fewer conserved syntenies with Hsa11, suggesting that they represent paralogons from earlier genome duplication events in stem vertebrates (VGD1 and VGD2). These phylogenetic and conserved synteny analyses converged to support the orthology of *APLNR* in human to *alpnr2* in zebrafish. This conclusion differs from an earlier report (Zhang et al., 2018) that did not consider phylogenetic analysis or analyze conserved synteny for the relevant parts of Hsa11 to Dre7. These considerations suggest that changing the name of the zebrafish gene *alpnr2* (ENSDARG00000004447) to *alpnr* would conform better to zebrafish nomenclature rules and better connect the zebrafish medical model to human biology.

Analysis of the zebrafish *aplnra* and *aplnrb* genes in the same tree (ENSGT01000000214406) showed that most teleost clades have orthologs of each gene, that the bonytongues, which branch deep in the teleosts, root each clade; furthermore, spotted gar and reedfish serve as pre-TGD outgroups, as expected from historical species relationships (Fig. 7B). The tree showed that the sister group of the ray-finned *aplnra+alpnrb* clade is a lobe-finned vertebrate clade rooted on coelacanth and amphibia as expected and includes ‘reptiles’ and birds (Fig. 7B). This clade, which we tentatively here call the *Aplnrl* clade, contains only monotremes and marsupials among mammals, indicating that this gene was lost in eutherian mammals (Zhang et al., 2018). Conserved syntenies showed that chicken (*Gallus callus*) chromosome Gga15, which contains *Aplnrl* (ENSGALG00000047717), has orthologs on both Dre8 and Dre10, the sites of *aplnra* and *aplnrb*, respectively, as well as Dre5 (Fig. 7D). These conserved syntenies independently verify that: 1) *Aplnrl* was present in the last common ancestor of human and zebrafish; 2) *Aplnrl* was lost from eutherians after they diverged from marsupials; and 3) the TGD produced *aplnra* and *aplnrb* paraogs.

These analyses support the hypothesis that the last common ancestor of zebrafish and human had at least two *Aplnr*-related genes; one became *APLNR* in human and ‘*aplnr2’* in teleosts and the other was lost in eutherians but retained in other tetrapods, and subsequently duplicated in the TGD, becoming ‘*aplnra’* and ‘*aplnrb’* in zebrafish. Renaming *aplnr2* (ENSDARG00000004447) to *aplnr, aplna* (ENSDARG00000002172) to *aplnrla*, and *aplnrb* (ENSDARG00000036670) to *aplnrlb* would better connect zebrafish to human biology.

Atlas cells expressing *aplnr* (ENSDARG00000004447) occupied midline fates, including prominently the floorplate (c176), hypochord (c218), and scleroderm (c33), as well as an embryonic intestinal epithelium cell type (c101). Expression of *alpnr* was detected in the spleen and heart of zebrafish adults by qPCR (Zhang et al., 2018) and we detected low expression in larval spleen cluster c162 but not in embryonic spleen cluster c210. Expression of *aplnr* in the Atlas was weak in three cardiac neural crest cells (c69) at 1dpf and a single cardiac muscle cell (c205) at 2dpf but was not expressed in the heart primordium (c147) at 1dpf or embryonic heart tube at 1 and 2dpf (c130). Expression of *aplnrla* was strongest in endothelial cell cluster c79, which arterial markers *flt4* and *ephb4* showed to represent arterial cells (c88, Fig. 7G); thus, venous cells were expressing the ligand *apln* (Fig. 7E) and arterial cells were expressing the receptor gene *aplnrla*. In addition, *aplnrla* was expressed in several clusters of basal skin cells (c46, c70, c108, c118, c145), in the pharyngeal endoderm (c45, c179), in the retina (c83) and lens (c106), confirming and extending previous reports (Helker et al., 2015; Kwon et al., 2016; Zhou et al., 2015). Expression of *aplnrlb* was detected in the Atlas both in the venous and arterial cell clusters (c79 and c88, Fig. 7H), as well as in a few slow muscle cells at 1dpf (c209) and several myoblasts at 2 and 5dpf (c84, c44), confirming *in situ* hybridizations (Tucker et al., 2007). These single cell gene expression clusters are consistent with Apelin signaling in COVID-19 comorbidities because they act in cardiovascular development (Deshwar et al., 2016; Scott et al., 2007; Zeng et al., 2007).

## DISCUSSION

These analyses show that: 1) nearly all components of the human Renin-Angiotensin-Aldosterone-System are conserved in zebrafish; 2) that zebrafish has multiple paralogs of some RAAS components and we identify their evolutionary relationships to human genes; 3) that zebrafish and human RAAS genes are generally expressed in equivalent cell types; and 4) that a specialized intestinal cell type is a focus of RAAS component expression including the SARS-CoV-2 receptor. Because many components of the RAAS function in zebrafish as in humans, these investigations focus attention on genes and cell types essential for utilizing zebrafish as a model to understand mechanisms leading to COVID-19 comorbidities.

### Zebrafish orthologs and co-orthologs of human RAAS components related to COVID-19 comorbidities

Key conserved RAAS features include *ace2* – the zebrafish ortholog encoding the SARS-CoV-2 receptor – and *ace* – the zebrafish ortholog encoding the enzyme that produces Ang II, which exacerbates COVID-19 comorbidities and likely worsens disease outcomes. Zebrafish has a functional Renin enzyme (Gomez et al., 1988; Hoshijima and Hirose, 2007; Jones et al., 1990; Liang et al., 2004) that acts on its structurally conserved Angiotensinogen protein (Lu et al., 2016a) to produce Angiotensin with a sequence that matches the ancestral tetrapod sequence and differs from the human form at three positions. After cleavage by Ace, zebrafish Ang II differs at position 1 and 5 from the human form. The zebrafish genome preserves enzymes that metabolize Angiotensin-related peptides, including importantly, Ace, Ace2, Enpep, and Anpep. Zebrafish conserves genes encoding Ang II receptors Agtr1 and Agtr2. Zebrafish also has orthologs encoding Slc6a19, which binds Ace2 and Adam17, the enzyme that creates the soluble form of Ace2. Furthermore, zebrafish has an ortholog encoding Tmprss2, which activates coronavirus spike protein for binding to ACE2 and bringing the virus into human cells. Zebrafish also has the ligand and receptors for the Apelin signaling system.

Zebrafish has a single ortholog of most RAAS genes but has duplicates of some. Many zebrafish duplicates of human RAAS genes derive from the teleost genome duplication event, including 1) *agtr1a* and *agtr1b*, 2) *anpepa* and *anpepb*, 3) *slc6a19a* and *slc6a19b*, and 4) *aplnrla* and *aplnrlb*. Other duplicated RAAS genes appeared by tandem duplication, including 1) *anpep* and *anpepl*, 2) *anpepla*.*1* and *anpepla*.*2*; and 3) *slc6a19a*.*1* and *slc6a19a*.*2*. The significance of these discoveries is that the actions of both zebrafish co-orthologs must be considered when translating zebrafish science to human biology.

The RAAS/Apelin system also provides an example of ‘ohnologs gone missing’, in which one ohnolog (paralog from a genome duplication) arising in vertebrate genome duplication events is lost in either ray-finned or lobe-finned vertebrates but reciprocally retained in the other taxon (Postlethwait, 2007); this situation happened for Apln receptor genes.

Two major RAAS components are missing from zebrafish. First, zebrafish and other ray-finned vertebrates, lack an ortholog of the Ang1-7 receptor *MAS1* (Bader et al., 2014; Fournier et al., 2012; Hoffmann et al., 2018). *MAS1* is a member of a large gene family derived by genome and tandem duplication events, followed by chromosome rearrangements (Rinne et al., 2019). Some other G-protein-coupled receptor may perform a similar function in ray-finned vertebrates or Ang1-7 may not function in ray-finned vertebrates.

The second component of the RAAS missing from zebrafish is aldosterone. In tetrapods, the adrenal steroid aldosterone acts on kidney collecting ducts to stimulate reabsorption of sodium and secretion of potassium and hydrogen ions, thereby regulating water balance and blood pressure. In contrast, teleosts, cartilaginous fishes, and jawless fishes lack aldosterone (Bridgham et al., 2006; Jiang et al., 1998; Nunez and Trant, 1999; Simpson and Wright, 1970). Aldosterone emerged in tetrapods after the evolution of the ability of Cyp11b1 to hydroxylate corticosterone and produce aldosterone (Bulow and Bernhardt, 2002; Jiang et al., 1998; Nonaka et al., 1995). Teleosts nevertheless have the aldosterone (mineralocorticoid) receptor Nr3c2 (Bridgham et al., 2006), which in fish is stimulated by either the aldosterone precursor 11-deoxycorticosterone or by cortisol (Kumai et al., 2014; Sturm et al., 2005). In the Atlas, *nr3c2* was expressed in embryonic periderm and larval gills, which help regulate salt balance at these developmental stages (Dymowska et al., 2012; Fu et al., 2010; Hoffmann et al., 2018; Rombough, 2007), suggesting that an ancestral role of Nr3c2 included salt and water balance even in the absence of aldosterone. Zebrafish mutants lacking active *nr3c2* responded poorly to a swirling stress test, but their response to hypotonic or hypertonic stress has not been reported (Faught and Vijayan, 2018). We also don’t know whether zebrafish Angiotensin receptors stimulate the release of 11-deoxycorticosterone, cortisol, or another steroid.

The conservation of nearly all RAAS components makes zebrafish an appropriate model to investigate the roles of the RAAS in the generation of COVID-19 comorbidities.

### Zebrafish express COVID-19-related RAAS genes in tissues similar to human

Results showed that, like humans and other mammals, zebrafish liver cells express *Angiotensinogen*. In the Atlas, 5dpf zebrafish larvae had three types of hepatocytes, two of which expressed *agt*, adding cellular precision to expression studies in adult zebrafish (Cheng et al., 2006). Induction of *Agt* expression in mammals relies on cortisol and inflammation (Brasier and Li, 1996; Demura et al., 2015). Cortisol binds the glucocorticoid receptor Nr3c1 and inflammation acts via interleukins and Tnfa to cause CCAAT-binding proteins like Cebpb and Cebpd to bind *Agt* enhancer elements. The co-expression of *cebpb, cebpd*, and to some extent *nr3c1* shows that gene expression is as expected if the initiation of the RAAS cascade is similar in humans and zebrafish.

Angiotensinogen is cleaved to form Ang I by Renin, which adult kidney juxtaglomerular cells secrete in mammals and zebrafish (Gomez et al., 1988; Hoshijima and Hirose, 2007; Jones et al., 1990; Liang et al., 2004)}(Rider et al., 2015). Renin-expressing cells in fetal mammals first appear in several tissues and organs, but predominantly in the adrenal (Gomez et al., 1988; Sequeira Lopez et al., 2004). Zebrafish apparently shares this developmental pattern because one of the three cells in the Atlas expressing *Renin* was a cell of the interrenal (the fish adrenal equivalent) (Liang et al., 2004), and the other two were cell types appropriate for different organs, as in mammals.

Ace, which cleaves Ang I to Ang II, was surprisingly shown by our scRNA-seq analysis to be co-expressed almost exclusively in a specific cell type in the zebrafish larval intestinal epithelium along with several other RAAS-components. The Atlas identified four different types of intestinal epithelial cells, one embryonic cluster and three larval clusters (Farnsworth et al., 2020). Larval intestinal cluster c152 expressed not only *ace*, but also *ace2*, the human ortholog of which encodes the enzyme that degrades Ang II and serves as a SARS-CoV-2 receptor. The coronavirus spike protein is activated after cleavage by Tmprss2, which stimulates the entry of the coronavirus into cells (Hoffmann et al., 2020; Millet and Whittaker, 2015; Walls et al., 2020). We found that *tmprss2* is also expressed in c152 and two other intestinal epithelium cell types in the Atlas, and it is also strongly expressed in the human digestive system (Fagerberg et al., 2014). Another protease, Ctsl, cleaves the spike protein of SARS-CoV (Simmons et al., 2005) and we show that *ctsla* is also strongly expressed in *ace2*-expressing c152 cells in zebrafish.

The c152 cell type also makes the enzymes that cleave Ang peptides encoded by *enpep, anpepa, anpepb*, and *dpp4*. Expression of these genes makes the human homolog of c152 cells a nexus of coronavirus infections because: 1) ANPEP is a receptor for the human common cold coronavirus HCoV-229E (Fehr and Perlman, 2015), 2) DPP4 is the receptor for MERS-CoV, the causative agent of the Middle East Respiratory Syndrome (Raj et al., 2013), 3) ACE2 is the receptor for SARS-CoV-2. This cell type is likely responsible for the digestive tract symptoms experienced by many COVID-19 patients.

C152 cells also express zebrafish *slc6a19* genes. In human, SLC6A19 binds ACE2 to form the SARS-CoV-19 receptor (Yan et al., 2020). The co-expression of *ace2* and *slc6a19* in zebrafish suggests functions shared with human. SLC6A20, which our genomic analysis shows is a likely tandem duplication-derived paralog of SLC6A19, also interacts functionally with ACE2 (Kristensen et al., 2011; Vuille-dit-Bille et al., 2015). Significantly, *SLC6A20* is at the peak of the strongest of two genome wide association study loci for undesirable COVID-19 outcomes (Ellinghaus et al., 2020). We suggest the hypothesis that genetic variants near *SLC6A20* affect the severity of COVID-19 symptoms due to variations in interactions with ACE2.

Angiotensin peptides are related to COVID-19 comorbidities because they bind to Agtr1 and Agtr2 receptors on vascular cells to help regulate vasoconstriction, and on the adrenal to stimulate secretion of aldosterone, leading to salt and water retention related to the comorbidity of obesity-related kidney damage (Fyhrquist and Saijonmaa, 2008). As in humans, zebrafish scRNA-seq analysis showed that *agtr1b* is expressed in endothelial cells and confirmed that *agtr2* is also expressed in endothelial cells (Wong et al., 2009).

The conserved expression of zebrafish RAAS-related genes supports the contention that RAAS regulation and function are similar in zebrafish and human and the novel finding of the nexus of RAAS component expression in a specific intestinal epithelial cell type focuses future research on c152 cells.

### The zebrafish RAAS functions like that of mammals

Hypertension, obesity, and diabetes are major COVID-19 comorbidities (Richardson et al., 2020) and all relate to the role of the RAAS in salt and water balance, vessel function, and inflammation, which all lie downstream of Ang II (Cabandugama et al., 2017; Fyhrquist and Saijonmaa, 2008). In humans, adrenal-derived glucocorticoids and inflammation induce Angiotensinogen (Brasier and Li, 1996; Stockhammer et al., 2010); inflammation also upregulates *agt* in zebrafish (Stockhammer et al., 2010), suggesting conserved regulatory mechanisms. The role of cortisol in *agt* regulation in zebrafish remains to be investigated.

As in human, *Renin* transcription in zebrafish is regulated by plasma salt concentration (Hoshijima and Hirose, 2007; Rider et al., 2015). In larval zebrafish, levels of the Renin product Ang II increase after exposure to acidic or ion-poor water (Kumai et al., 2014) similar to the response of Ang II in humans with high levels of circulating sodium (Crowley and Coffman, 2012). Zebrafish *ren* mutants are normally adult viable but have enlarged swim bladders, expected from aberrant water balance, and dramatically more Renin cells in the kidney, suggesting regulation by a negative feedback loop (Mullins et al., 2019). Treating larval zebrafish with fish Ang I or Ang II peptides caused animals to accumulate Na^+^ (Kumai et al., 2014), showing that zebrafish larvae respond as expected from the mammalian system. In mammals, Ang II acts on sodium uptake via aldosterone binding to the aldosterone receptor, but fish have no aldosterone, so although the input and outcome are similar between these taxa, the hormones must differ; likely either cortisol or 11-deoxycorticosterone in fish (Sturm et al., 2005). Recent investigations in zebrafish with knockout mutations in the glucocorticoid receptor and the mineralocorticoid receptor promise to clarify these issues (Faught and Vijayan, 2018; Faught and Vijayan, 2020; Ziv et al., 2013).

The zebrafish RAAS is pharmacologically similar to that of mammals. The Ace inhibitor lisinopril blocks the effects of Ang I on sodium uptake in zebrafish, as predicted if Ace were required to convert Ang I to Ang II (Kumai et al., 2014). Zebrafish cultured from 24hpf to 96dpf in water containing the Ace inhibitor captopril do not differ in survival from controls and neither do fish in water with 5% of the normal salt concentration (Rider et al., 2015), but the combination of captopril and low ionic strength water reduces the survival of larval zebrafish by about 95% and causes an increase in Renin expression (Rider et al., 2015). Ace inhibition by captopril also upregulates *ren* in adult and larval zebrafish (Hoshijima and Hirose, 2007; Rider et al., 2017; Rider et al., 2015) and affects cardiovascular function (Margiotta-Casaluci et al., 2019). The Ace inhibitor enalapril, used to treat patients with high blood pressure, causes intraocular blood vessels to dilate in zebrafish (Kitambi et al., 2009) as expected from conserved functions. The fact that several different Ace inhibitors have similar effects in humans and zebrafish supports both the physiological conservation of the Ace (Prokop et al., 2015).

Drugs that block the Ang II receptor Agtr1 also provide evidence that the RAAS works similar in zebrafish and human. The selective Agtr1 antagonist telmisartan blocks sodium uptake in zebrafish cultured in low ionic strength water as expected if Ang II acts on sodium retention by binding to Agtr1 (Kumai et al., 2014), even though fish lack aldosterone. The high blood pressure medication fimasartan, an Ang II receptor antagonist, also ameliorates zebrafish models of heart failure, normalizing the expression of atrial natriuretic peptide, reducing cell death around the heart, and improving blood flow (Quan et al., 2020). These results show that Agtr1 structure and function are conserved in human and zebrafish.

### Conclusions

These discussions show that zebrafish has nearly all RAAS and Apelin signaling components, distinguish between previously confusing zebrafish orthologs and paralogs of human RAAS-related genes, identify for the first time specific cell types in which zebrafish RAAS components are expressed, and first identify a specific intestinal epithelial cell type that is a previously unrecognized focus of RAAS component expression in zebrafish and likely other vertebrates, including humans. Coupled with data from the literature showing that the RAAS functions in similar ways in zebrafish and human and that RAAS-related drugs tend to act in zebrafish as they do in humans makes zebrafish an ideal model for interrogating the roles of the RAAS in the origin of COVID-19 comorbidities. Coupled with the exquisite imaging that zebrafish embryos and larvae provide and the ease of precise mutagenesis, zebrafish offers a highly useful system for research into the mechanisms of COVID-19 pathologies and comorbidities.

## MATERIALS AND METHODS

### Phylogenies and conserved syntenies

To identify genes that encode zebrafish proteins related to the human RAAS, we utilized phylogenetic analysis embedded in the ComparaTree method at Ensembl (Flicek et al., 2010). This method uses phylogenetic distance and tree structure to identify gene duplications. To verify orthologs and co-orthologs of mammalian genes, we analyzed conserved syntenies using the Synteny Database (Catchen et al., 2009). Nomenclature rules for genes and proteins in human, mouse, and zebrafish use conventions described at the Zebrafish Information Network (ZFIN, https://wiki.zfin.org/display/general/ZFIN+Zebrafish+Nomenclature+Conventions). For non-human sarcopterygians, we use mouse conventions and for actinopterygians, we use zebrafish conventions.

### Single cell transcriptomics

Expression patterns of zebrafish RAAS components were identified from the zebrafish single cell transcriptomic Atlas (Farnsworth et al., 2020).

### Animals

Zebrafish (*Danio rerio*) were reared using standard husbandry ((Westerfield, 2007). Strains used were *Tg(olig2:GFP)vu12* for two samples at 1 and 5dpf and one sample at 2 dpf, and *Tg(elavl3:GCaMP6s)* for one sample at 2dpf. Embryos were dissociated as described (Farnsworth et al., 2020). Fifteen animals of mixed but unknown sex were used for each of the six batches. Work was conducted under the approval of the University of Oregon IACUC, Protocol # 18-31.

### Reagents

Dissociated cells were prepared for scRNA-seq using the 10X Chromium platform v.2 chemistry. cDNA libraries were amplified by 15 PCR cycles and sequenced on Illumina Hi-seq or Next-seq.

### Statistics

Sequencing data were analyzed by Cell Ranger version 2.2.0 (Zheng et al., 2017) and Seurat (Satija et al., 2015). For further details, see (Farnsworth et al., 2020) and https://www.adammillerlab.com/.

### Data availability

scRNA-seq data are publicly available at https://cells.ucsc.edu/?ds=zebrafish-dev.

## ACKNOWLEDGEMENTS

We wish to thank the University of Oregon Aquatic Animal Care Services director T. Mason for fish care and M. Haeussler and M. Speir for hosting data on the UCSC Cell Browser.

## COMPETING INTERESTS

Authors declare no competing or financial interests.

## FUNDING

Funding was provided by the National Institutes of Health Office of the Director grant R24OD026591 (JHP and ACM) and the University of Oregon (ACM).

## DATA AVAILABILITY

All scRNA-seq data are publicly available at https://cells.ucsc.edu/?ds=zebrafish-dev.

## AUTHOR CONTRIBUTIONS STATEMENT

Conceptualization and writing (JHP); Reviewing and editing (ADM, DRF); Designing methodology (JHP, ACM, DRF); Developing, implementing, and maintaining computer code (ACM); Funding acquisition (JHP, ADM); Performing experiments and collecting data (JHP, DRF, ADM).

## FIGURE LEGENDS

**Figure 1**. ***Angiotensinogin* and *Renin***. A. The section of *Homo sapiens* chromosome 1 (Hsa1) that contains *AGT* is conserved with the segment of *Danio rerio* chromosome 13 (Dre13) that contains *agt*. B. The 220 clusters from the zebrafish scRNA-seq Atlas with liver clusters boxed (Farnsworth et al., 2020). C. Cells in hepatocyte cluster c217 and a sub-set of hepatocytes in c121 express *agt*. Each dot represents a cell. Red color intensity indicates expression level. Blue cells are not expressing. D. Expression of the Agt-regulator *cebpb* in larval liver cells. E. Conserved syntenies confirm orthology of zebrafish *ren* to *REN*. F-G. *ren* (orange cell) was expressed in c87, part of which contains precursors to the interrenal, the fish equivalent of the adrenal cortex, as marked by expression (circled cells) of cortisol-synthesis genes *star* (F) and *hsd3b1* (G).

**Figure 2**. **Evolution of Angiotensin sequences**. Arrows indicate human and zebrafish sequences. Species organized according to published phylogenies (Hughes et al., 2018; Upham et al., 2019). Ang I and Ang II indicated at far right. For Latin names and accession numbers, see Supplementary Table S1.

**Figure 3**. **Genomics and expression of *ace***, ***agtr***, **and *nr3c2***. A. Conserved syntenies verify orthology of zebrafish *ace* to human *ACE*. B. Expression of *ace* in a specific intestinal epithelial cell type in c152. C. Clusters c101, c152, c168, and c214 are intestinal epithelial cell types (Farnsworth et al., 2020). D. The age of each cell, as indicated in the insert, showed that *ace* is expressed in a 5dpf intestinal cell type c152 that does not exist at 1dpf or 2dpf. E. Double conserved synteny of zebrafish *agtr1a* and *agtr1b* to human *AGTR1*. F. *agtr1b* was expressed only in endothelial cells (c88). Expression of *agtr1a* was not detected. G. Conserved synteny of *agtr2* and *AGTR2*. H. Expression of *agtr2* was detected in c88 endothelial cells, including some in which *agtr1b* was detected. I. Conserved syntenies for the aldosterone receptor gene *nr3c2*. J. Expression detected for *nr3c2* in intestinal epithelial clusters c168 and c101. K.-L. Expression of *nr3c2* in the embryonic periderm and in larval gills.

**Figure 4**. **Genomics and expression of *ace2***, ***enpep***, ***anpep* and *dpp4***. A. Conserved synteny supports the orthology of human *ACE2* to zebrafish *ace2*. Expression of *ace2* (B), *enpep* (C), *anpeplb* (D), *anpepb*.*1*, (E), *anpepa* (F), and *dpp4* (G). All were expressed in c152, the cell type expressing *ace*, except *anpeplb*, which was expressed in larval blood vessels and a few neural crest cells, but not in intestinal epithelium.

**Figure 5**. **Conserved syntenies and expression of *SLC6A19*-related genes**. A. Conserved syntenies showing paralogy of a part of Dre16 containing *slc6a19b* and a portion of Dre19 containing *slc6a19a*.*1* and *slc6a19*.*2* and their relationship to human chromosome Hsa5 around *SLC6A19*. B-D. Expression of *slc6a19a*.*1, slc6a19a*.*2*, and *slc6a19b*, respectively, in the zebrafish Atlas in c152 and c168, which represent the same larval intestinal epithelial cells that expressed *ace* and *ace2*.

**Figure 6**. **Conserved syntenies and expression of a*dam17***, ***tmprss2***, **and c*tsl***. A. Double conserved synteny between Dre17 and Dre20 confirming co-orthology of *adam17a* and *adam17b* to *ADAM17* and their origin in the TGD. B. Expression of *adam17a* weakly in intestinal epithelium cells and (C) stronger in the vascular endothelium. D. Conserved syntenies verify orthology of *tmprss2* to *TMPRSS2*. E. Expression of *tmprss2* was detected in one cell in intestinal epithelium c168 and in 12 other cells broadly dispersed in the atlas. F. Human *CTSL* shares conserved syntenies with zebrafish *ctsla*. G, H. Expression of *ctsla* in the endothelium and intestinal epithelium.

**Figure 7**. **Apelin and its receptors**. A. Phylogenetic tree (ENSGT01000000214406) with the *APLNR/aplnr2* (or *aplnr* (ENSDARG00000004447) subtree (blue) expanded. B. The same tree with the *aplnra/aplnrb* (or *aplnrla*, ENSDARG00000002172) / *aplnrlb* (ENSDARG00000036670)) subtree (orange) expanded. C. A dot plot representing orthologs and paralogs of Hsa11 genes vs. zebrafish (Dre) chromosomes plotted directly above the location of Hsa11 genes showing extensive conservation with Dre7, the site of *aplnr*, but not Dre8 or Dre10, the locations of *aplnrla* and *aplnrlb*, respectively. D. A dot plot of chicken chromosome Gga15 vs. zebrafish chromosomes showing the sites of *aplnrla* and *aplnrlb* and their ortholog in chicken (*Aplnrl*, ENSGALG00000047717) identified in the tree in panel B. E.-H. Expression of *apln, aplnr2, aplnra*, and *aplnrb* in endothelial cells.

